# neoMS: Attention-based Prediction of MHC-I Epitope Presentation

**DOI:** 10.1101/2022.05.13.491845

**Authors:** Nil Adell Mill, Cedric Bogaert, Wim van Criekinge, Bruno Fant

## Abstract

Personalised immunotherapy aims to (re-)activate the immune system of a given patient against its tumour. It relies extensively on the ability of tumour-derived neoantigens to trigger a T-cell immune reaction able to recognise and kill the tumour cells expressing them. Since only peptides presented on the cell surface can be immunogenic, the prediction of neoantigen presentation is a crucial step of any discovery pipeline. Limiting neoantigen presentation to MHC binding fails to take into account all other steps of the presentation machinery and therefore to assess the true potential clinical benefit of a given epitope. Indeed, research has uncovered that merely 5% of predicted tumour-derived MHC-bound peptides is actually presented on the cell surface, demonstrating that affinity-based approaches fall short from isolating truly actionable neoantigens. Here, we present neoMS, a MHC-I presentation prediction algorithm leveraging mass spectrometry-derived MHC ligandomic data to better isolate presented antigens from potentially very large sets. The neoMS model is a transformer-based, peptide-sequence-to-HLA-sequence neural network algorithm, trained on 386,647 epitopes detected in the ligandomes of 92 HLA-monoallelic datasets and 66 patient-derived HLA-multiallelic datasets. It leverages attention mechanisms in which the most relevant parts of both putative epitope and HLA alleles are isolated. This results in a positive predictive value of 0.61 at a recall of 40% on its patient-derived test dataset, considerably outperforming current alternatives. Predictions made by neoMS correlate with peptide identification confidence in mass spectrometry experiments and reliably identify binding motif preferences of individual HLA alleles thereby further consolidating the biological relevance of the model. Additionally, neoMS displays extrapolation capabilities, showing good predictive power for presentation by HLA alleles not present in its training dataset. Finally, it was found that neoMS results can help refine predictions of response to immune checkpoint inhibitor treatment in certain cancer indications. Taken together, these results establish neoMS as a considerable step forward in high-specificity isolation of clinically actionable antigens for immunotherapies.

## Introduction

Since the approval of immune checkpoint inhibitors (ICIs) for clinical use across multiple indications, immunotherapy has grown into an essential strategy in the fight against cancer. The main actors driving an anti-tumour response subsequent to ICI treatment, are neoantigen-specific T-cells, cognate to peptides uniquely presented on tumour cells due to tumour-specific genetic alterations (Miao et al., 2018; Rizvi et al., 2015; Snyder et al., 2014). Consequently, therapies tailored to target patient-specific neoantigens represent a particularly promising avenue to deliver additional clinical benefit to patients. Such personalised immunotherapies include therapeutic vaccines, which aim at priming and boosting T-cell responses towards a specific set of neoantigens, and adoptive cellular therapies involving T-cells targeted against the tumour which are grown, stimulated and potentially engineered *ex vivo* before being (re)introduced into the patient(Waldman et al., 2020).

These tailored approaches rely on the ability to correctly identify tumour-derived neoantigens likely to elicit a T-cell immune reaction. For neoantigens to be able to prime CD8 T-cells, presentation at the tumour cell surface by the class I major histocompatibility complex (MHC-I) is required, thus the prediction of peptide-MHC (pMHC) presentation is a crucial step of any neoantigen discovery pipeline. Most state-of-the-art pipelines only use the predicted binding affinity of a given peptide to one or several HLA alleles as a proxy for overall presentation likelihood and by extension immunogenicity (P. Ott et al., 2017; P. A. Ott et al., 2020; Sahin et al., 2017). Indeed, deep learning models such as netMHC, netMHCpan and MHCflurry achieve excellent performance as predictors of binding affinity(Jurtz et al., 2017; O’Donnell et al., 2018; Zhao & Sher, 2018). These models however do not take into account many additional steps in the epitope presentation process such as protein degradation by the proteasome, peptide loading complex processing, and peptide-MHC stability (Chong et al., 2018; Kincaid et al., 2016). This results in rather poor prediction of actual presentation at the cell surface (Bulik-Sullivan et al., 2019).It has been shown that as little as 5% of tumour-derived peptides predicted by the models to bind the MHC are actually presented (Bassani-Sternberg et al., 2015; Yadav et al., 2014), demonstrating the insufficiency of affinity-based approaches to isolate actionable neoantigens. This lack of specificity can have major practical consequences on the clinical development of neoantigen-based therapies. A recent consortium study showed that only 6% of neoantigens prioritised on binding affinity among other parameters could actually elicit an immune response in 6 patients with either melanoma or non-small cell lung cancer. This highlights both the importance of epitope presentation as a selection criterion and the shortcomings of current prediction methods(Wells et al., 2020).

Based on these findings, binding affinity-trained models are often combined in a prediction workflow with other tools trained separately on specific data, such as *e.g. in vitro* proteasome digestion data, to predict other steps of the antigen presentation pathway(Calis et al., 2015). Examples of such tools are NetChop20S, NetChopCterm, and ProteaSMM for MHC class I compatible peptide proteolysis(Nielsen et al., 2005; Tenzer et al., 2005), with NetChopCterm showing the most accurate predictions. Furthermore, PRED^TAP^ and the support-vector-machine based TAPREG can be used for TAP associated transport simulations(Diez-Rivero et al., 2010; Zhang et al., 2006). However, due to the limited size of their training datasets, these tools often suffer from stronger predictive inaccuracies than pMHC binding affinity predictions. As these imprecisions get amplified and compounded in a convoluted pipeline, the overall added predictive value often fails to overcome the increase in noise.

the availability of mass spectrometry (MS)-derived MHC ligandomic data has opened the door to the development of a new generation of presentation prediction algorithms(Bulik-Sullivan et al., 2019; Chen et al., 2019; Hu et al., 2019; Sarkizova et al., 2019; Venkatesh et al., 2020). Some of these approaches involve monoallelic predictors where peptide presentation likelihood is predicted for single HLA alleles(Hu et al., 2019; Sarkizova et al., 2019; Venkatesh et al., 2020). In contrast, multiallelic predictors allow presentation prediction for an actual, patient-representative HLA allele set. In the strategy displaying the best results, a deep-learning model is built for each HLA allele and the presentation prediction of a given peptide by an actual set of HLA alleles is derived by summing the presentation prediction across every individual allele(Bulik-Sullivan et al., 2019). These methods can achieve performances significantly better than the affinity-based models, with a maximum positive predictive value (PPV) of 0.45 at a recall rate of 40% for presentation predictions in clinically relevant samples, expressing up to 6 different HLA-I alleles(Bulik-Sullivan et al., 2019).

While these models represent a considerable improvement over previous predictors, they still have notable drawbacks for clinical applicability. Although they are able to extrapolate and produce predictions for individual alleles not present in their training set, monoallelic predictors work on a one-to-one basis, and fail to correct for potential interactions between the processing pathways of different HLA alleles likely to happen in multiallelic cells. Conversely, current multiallelic predictors rely on training data availability, as they can only output predictions for HLA sets at least partially seen during training; however, they are more representative of the biological reality. In addition, both approaches do not leverage the recent progresses made in the field of deep learning. Most notably, since the epitope presentation problem aims at extracting information from pairs of sequences (the epitope amino-acid sequence and the HLA sequence), it can be approximated to a natural language processing (NLP) problem. In this respect, one of the most important recent breakthroughs in NLP models has been the inclusion of attention mechanisms (Vaswani et al., 2017) to prioritise and infer relationships between words in a sentence, thus forming a composite representation of language. Several deep learning architectures, such as the Transformer and the slightly modified BERT-like models have successfully used attention to achieve remarkable performances in NLP tasks such as language inference and text mining(Devlin et al., 2019; Lee et al., 2020; Martin et al., 2019). Furthermore, these architectures have already been adapted to tackle biological questions like protein structural modeling, RNA-protein interaction, protein sequence evolution and even MHC binding affinity prediction, often setting new performance records(Gao et al., 2020; Rives et al., 2000; Venkatesh et al., 2020; Yuan & Yang, 2021).

Here, we present neoMS, an attention-based deep learning algorithm for MHC-I epitope presentation prediction in clinically relevant (ie, HLA-multiallelic) settings. The attention mechanism is used to identify and prioritise relevant amino acid subsequences within both epitope and HLA allele that would most inform the presentation likelihood. Trained on 386,647 epitopes detected in the ligandome of 92 unique HLA alleles and 66 multiallelic datasets, neoMS achieves a PPV of 0.61 at a recall of 40% on a test set of clinically relevant tumour samples, and significantly outperforms current state-of-the-art presentation prediction methods. Binding motif preferences of individual HLA alleles can reliably be identified by neoMS, and its predictions correlate well with peptide identification confidence levels in out-of-training MS datasets, highlighting the biological relevance of the model. We also show that neoMS is capable of extrapolation, showing good predictive power for alleles not present in the training set. Finally, we demonstrate that the amount and rate of neoantigen presentation as predicted by neoMS correlates with response to ICI treatment in several cancer indications, and that the neoMS output can be combined with well established metrics such as tumour mutational burden (TMB) to further refine ICI response prediction models in certain indications.

## Results

### neoMS adopts a transformer-like architecture and is trained on MS-derived data

Positive ligandomic data was obtained from publicly available sources(Abelin et al., 2017; Bulik-Sullivan et al., 2019; di Marco et al., 2017; Sarkizova et al., 2019; Trolle et al., 2016); we selected unique peptides of length 8-11 detected with high confidence (false detection rate, or FDR <0.02) by MS either in the ligandome of a monoallelic cell line or in the ligandome of normal or tumour multiallelic samples. This length range was chosen since it covers most previously identified presented MHC-I epitopes (Trolle et al., 2016). Our final dataset comprised 386,647 positive datapoints, of which 173,361 were derived from multiallelic HLA sets (comprising 92 different ligandomes) and 210,760 from monoallelic HLAs (66 ligandomes), and was split into three subsets in our initial setup. For all subsets, negative data was synthesised *in silico* by random selection of 8-11mers from the proteome, randomly attributing them to one of the HLA alleles or HLA sets present in the positive dataset counterpart. To generate our test dataset, 729 of the 386,647 positive instances, derived from five different tumour multiallelic samples were set aside, and 1,822,500 negative instances were synthesised with similar HLA makeup (thus reaching a 1:2500 ratio between positive and negative instances). This specific testing dataset was chosen for comparison purposes with other models (Bulik-Sullivan et al., 2019), and because it allowed to evaluate peptide presentation in the most clinically relevant set-up. Another 326,297 positive instances were used for training, along with 652,594 synthesised negative instances (1:2 ratio). The remaining 57,582 positive instances, along with 115,164 negative ones, were used as a validation set for model monitoring and early stopping during training

The neoMS model follows a BERT-like architecture (**Fig.1**), *i.e*. a bidirectional transformer architecture. Peptides and HLA alleles are ingested as a concatenated string (a single “sentence”) and tokenised at the character level, using individual amino acids as tokens. Membership to each component (*i.e*. either peptide or HLA) was provided by the model using a typing mask. Every instance started with a specific classification [CLS] token and sentences were separated by a [SEP] token. During the forward pass, tokenised sentences were embedded and then processed by four self-attention layers, each containing six attention heads; since every instance now consisted in a concatenated pair of sentences, this process effectively included bidirectional cross-attention between two sentences. The pooled attention outputs of the final layer hidden state towards the classification token was finally processed by a standard feed-forward classifier neural network to produce a probability of presentation. The neoMS model was trained for 6 epochs as performance on the validation set showed only minimal increases past that point. Model hyperparameters were finetuned to improve the chosen performance metrics, precision and recall.

**Fig. 1:**
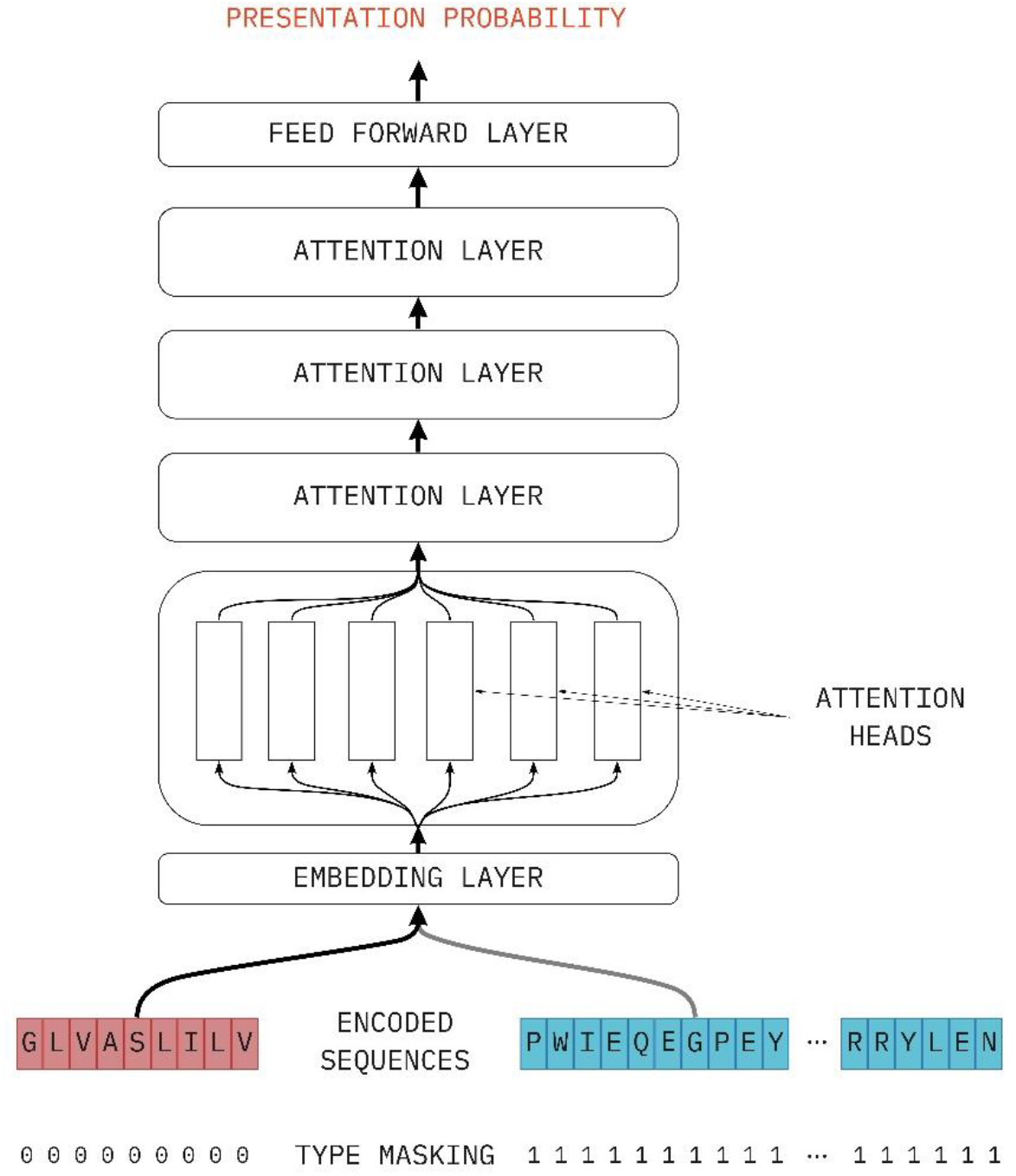
Schematic representation of the neoMS model Attention heads are shown only for the first attention layer.

### The neoMS algorithm outperforms current state -of-the art models on both in- and out-of-training HLA alleles

When confronted to the test dataset, the neoMS algorithm reaches a PPV of 0.61 at recall 40%. To assess its performance against other algorithms, we chose to compare it to the MS-only EDGE algorithm(Bulik-Sullivan et al., 2019), since the suite of EDGE model currently showcase the best performances on HLA-multiallelic datasets. While the full EDGE model achieves better performance than its MS-only counterpart, it uses additional patient-specific inputs that are out of the scope of the generalized framework that neoMS offers (such as expression level of the peptide), and is therefore not a suitable comparison. On the same test dataset, neoMS outperforms the current state-of-the art EDGE algorithm across the entire recall range (**Fig. 2A**). It also performs considerably better than affinity-based methods on the same dataset: the *MHCflurry* algorithm for example performs significantly worse, only reaching a PPV of 0.18 on average across all ranges. The poor performance of MHCflurry is principally due to the high false positive rate of the algorithm, as an important fraction of negative peptides are considered binding across all recall ranges.

**Fig. 2:**
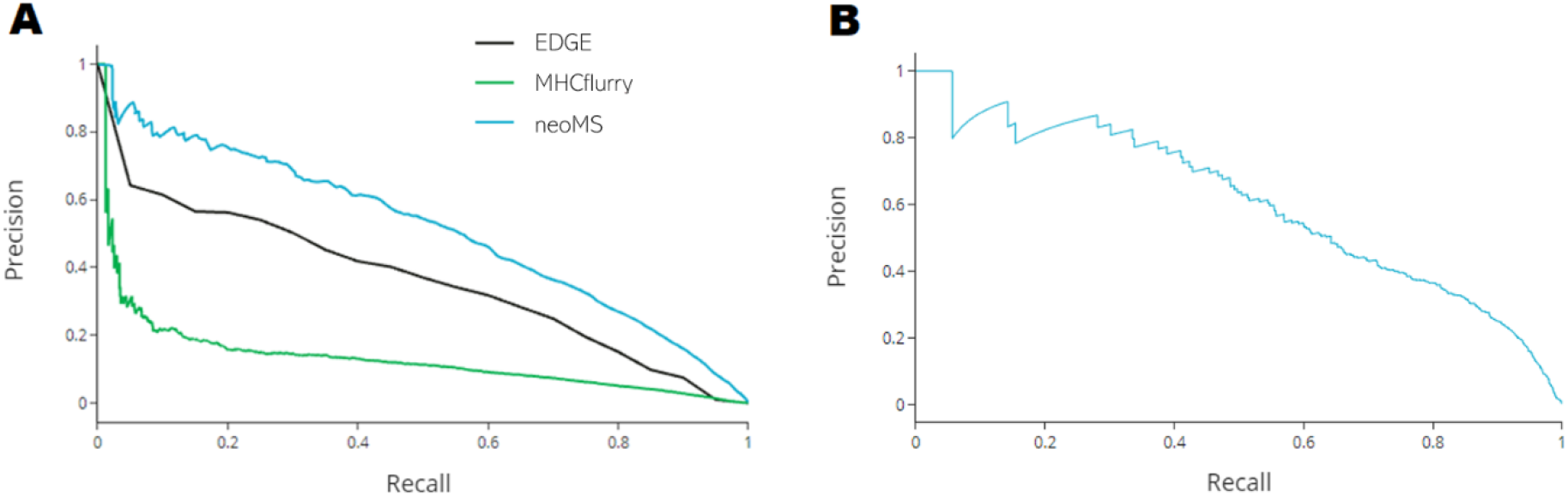
Performance of the neoMS model A. Test dataset precision-recall curves. The neoMS algorithm is shown in blue, current state-of-the-art model EDGE in black, and the affiniy-based model MHCflurry in green. B. A*74:01 dataset precision-recall curve of neoMS

Furthermore, one of the unique strengths of the neoMS architecture is its extrapolation capabilities, only shared by few predictors (Sarkizova et al., 2019). As a sequence-to-sequence prediction model, it is potentially able to assess presentation likelihood for alleles previously unseen in training, by using ligandomic information of training HLA alleles with similar sequences. To test this, we constructed a test set of 2039 peptides identified by MS in the ligandome of HLA-A*74:01, supplemented by synthesised negative instances at a ratio of 2,500 to 1. None of the training instances, be it mono- or multi-allelic, contained HLA-A*74:01, and no HLA allele present in the training set had the exact same sequence as HLA-A*74:01 (the closest one differing by 2 amino acids). As **Fig.2B** shows, neoMS displayed good prediction capabilities for A*74:01, thus demonstrating the extrapolation capabilities of the model.

### neoMS predictions correlate with identification confidence in MS experiments

While MS-based ligandomic analyses can achieve remarkable depth during peptide identification, inherent sources of background noise can lead to degrees of uncertainty in the peptides they identify. High confidence peptides display clear spectra, are supported by several scan instances, and are expected to be the most representative of the actual ligandome (*ie* to be “true positives”). As such, their presentation likelihood as scored by neoMS should be higher than the likelihood of their low-confidence counterparts.

To test this hypothesis, all peptides identified in the ligandomes of four publicly available hepatocellular carcinoma samples (Loffler et al., 2018) were binned in four ordinal categories reflecting their level of confidence, and their presentation likelihood was evaluated using neoMS. It is to be noted that this data had not been used in the training of neoMS. As shown in **Fig.3**, peptide identification confidence was strongly correlated with neoMS predictions, with high-confidence peptides consistently displaying higher neoMS scores across all samples tested (Kruskal-Wallis H test, p < 10^-200^, and Dunn’s *post-hoc* tests, p < 10^-4^). This confirms that the neoMS algorithm is able to correctly discriminate between high-confidence ligandomic peptides and potential noise.

**Fig.3:**
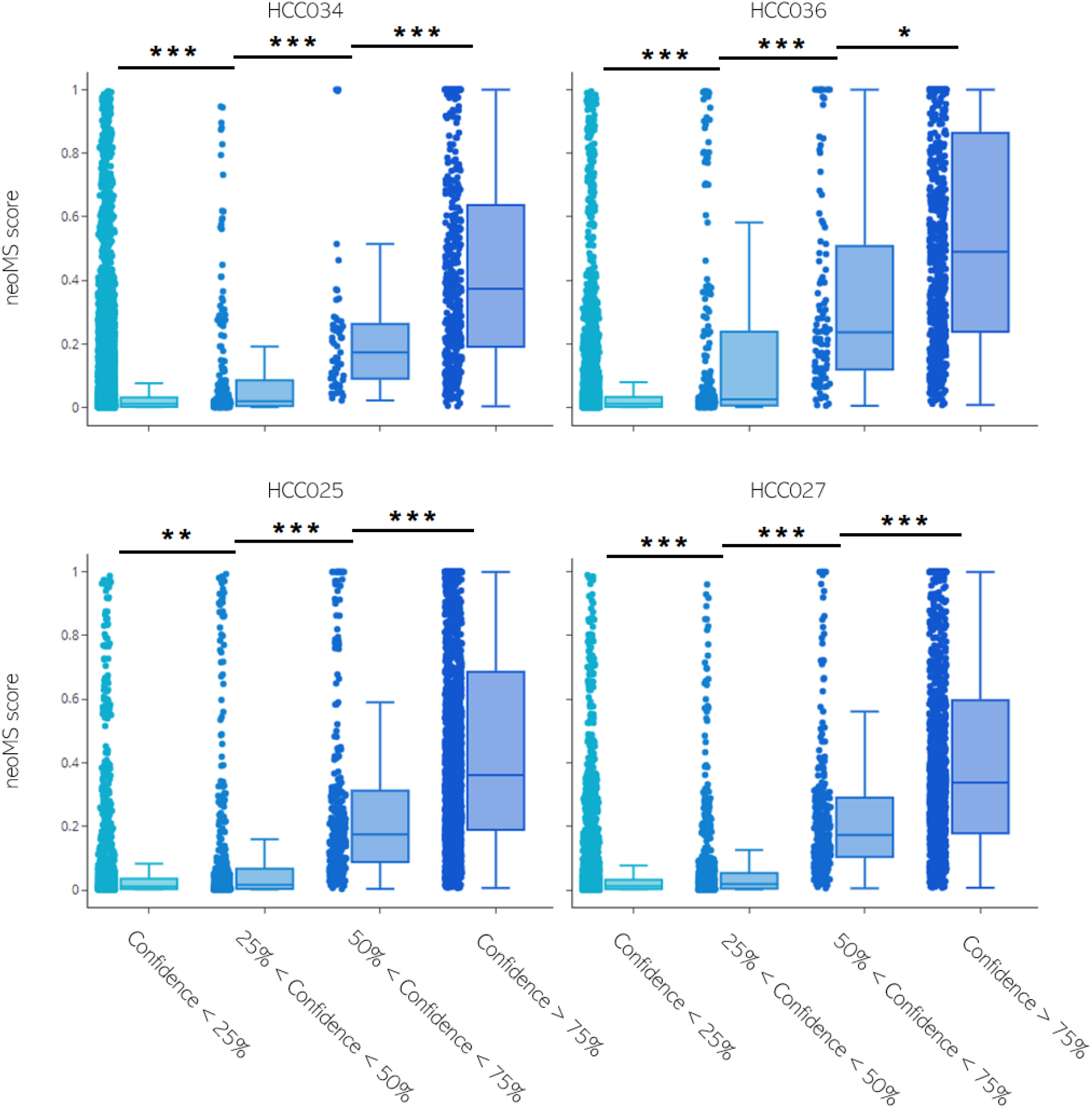
neoMS prediction scores as a function of peptide identification confidence level in mass spectrometry-derived ligandome experiments Each panel represents a different hepatocellular carcinoma sample. Each point represents a neoMS prediction for a given peptide; peptides were binned in quartiles according to their PeptideShaker confidence score. Stars indicate significant p-values for a Dunn’s post-hoc test following a statistically significant Kruskal-Wallis H-test. *, p<0.05; **, p<10^3^, ***; p<10^7^

### neoMS predictions can recapitulate HLA binding motifs

To assess the biological relevance of neoMS, we evaluated whether or not the peptide motifs learned during training reflected the key elements of the actual HLA binding motifs. By ranking neoMS predictions of a random set of peptides on monoallelic predictions and extracting the motifs of the top 1% presented peptides, we were able to recapitulate the binding motifs of HLA-A*02:01 (**Fig.4,** (Bassani-Sternberg et al., 2017; Solleder et al., 2020)), therefore establishing the biological relevance of the model.

**Fig. 4:**
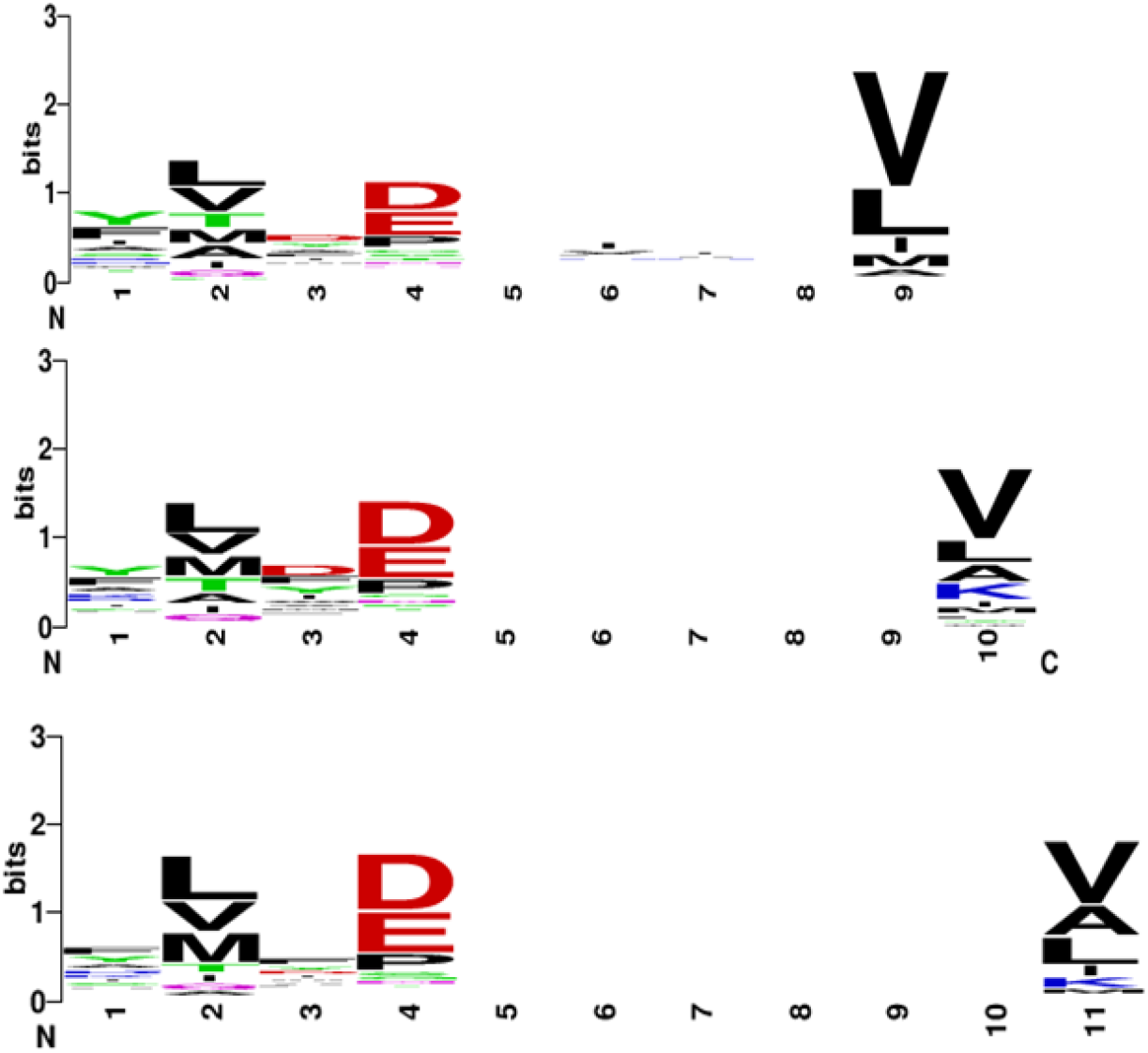
HLA*A02:01 preferred binding motifs isolated by neoMS.

### Attention mechanisms are a key process of the neoMS model

The key mechanism underlying the neoMS algorithm is the attention process, in which privileged relationships are established between sometimes distant amino acids both within each sequence considered and across sequences. This process can be visualised to examine which positions are most relevant to the presentation problem. As an example, the attention process occurring upon presentation prediction of the true positive GLVASLILV peptide by HLA-A*02:01 is illustrated in **Fig.5**. In each layer of the neoMS model (see **Fig.5B** for a full view of attention in the first layer of the model), specific relationships are established between subsequences of both peptide or HLA. Remarkably, these privileged relationships can span both the peptide and the HLA sequence, making it possible to identify complex interactions where several part of the core HLA sequences interact in synergy with one or more positions in the potential epitope. In one head of the first layer of neoMS, for instance (**Fig.5A**, marked position), position 33 of the first part of the core HLA sequence attends to both a specific position in the epitope, and to a position in the second part of the core sequence. Self-attention within each sequence considered by the model reveals intra-sequence dependencies: as demonstrated by the high level of attention given to them, it is possible to confirm for instance the importance of positions 2 and 9 of a peptide in predicting the likelihood of its presentation by a specific allele (**Fig.5D**). This importance is confirmed when considering cross-attention, in which focused relationships are established between the HLA sequence and the same positions (**Fig.5C**)

**Fig. 5:**
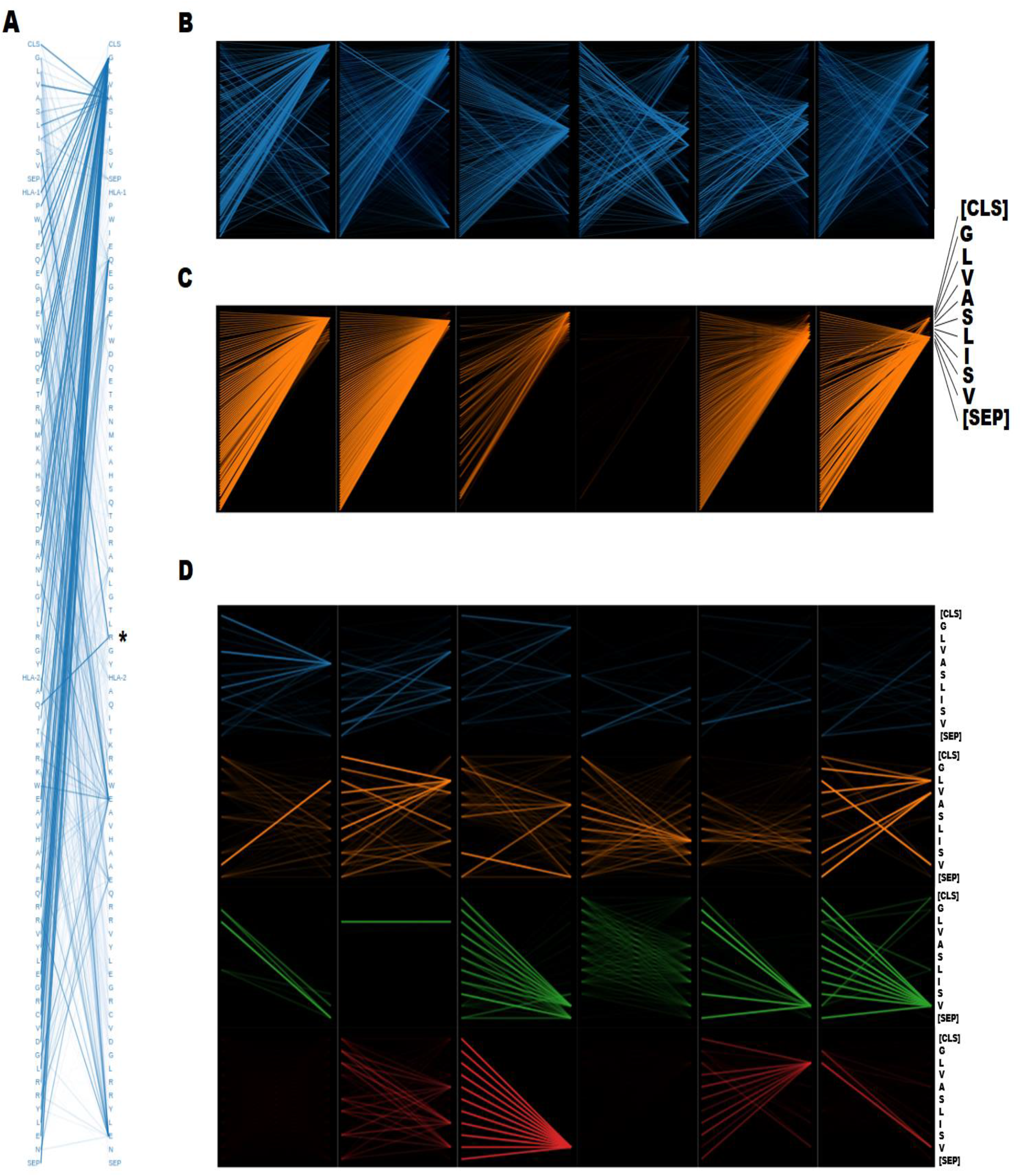
Visualisation of the attention mechanism in neoMS during presentation prediction The example being shown is the prediction of the presentation of the GLVASLISV by HLA-A*02:01 Thicker and brighter lines indicate higher levels of attention between a pair of amino-acids. Special tokens are denoted as follows: [CLS], special token marking the beginning of the sequence, [SEP], separator token between different input types, [HLA-1] and [HLA-2] tokens prefacing the first and second part of the HLA core sequence, respectively. A. Cross- and self-attention in the first head of the first attention layer of neoMS. The amino-acid marked with a star (*) demonstrates an example of attention across peptide and HLA sequences. B. Full cross- and self-attention of the first attention layer of neoMS. Each panel represents a different attention head. C. Attention of the HLA sequence to the epitope sequence in the second attention layer of neoMS. Each panel represent a different attention head. The peptide sequence being attended to (including special tokens) is shown on the right. D. Complete self-attention in neoMS for the epitope sequence. Each line of panel represents a different attention layer, from the first at the top to the last on the bottom. Individual panels on each line show individual attention heads. The epitope sequence (identical on both sides of each panel for self-attention) is shown on the right.

### neoMS can help refine response to ICI treatment prediction

Finally, the neoantigen presentation landscape characterized by neoMS was investigated in patient cohorts of two cancer indications. Since the neoantigen load derived from tumor-specific mutations is one of the main forces driving the antitumoural immune response unlocked by ICIs, it can be hypothesised that tumours responding to ICI treatment will exhibit a higher number of presented peptides as predicted by neoMS. Mutation data as well as clinical information was gathered from two publicly available cohorts - a set of 34 non-small cell lung carcinomas (NSCLCs) treated with Prembolizumab and a set of 64 melanomas treated with Ipilimumab (Rizvi et al., 2015; Snyder et al., 2014). The neoantigen landscape of each tumor was then computed and the number of presented neoantigens was predicted using neoMS.

As described previously(Rizvi et al., 2015; Snyder et al., 2014), tumours with high TMB were more likely to respond favourably to ICI treatment regardless of the cancer indication (Fig.6A,D, Mann-Whitney test, p = 0.02 for the NSCLC cohort and p = 0.01 for the melanoma cohort). The number of presented neoantigens as predicted by neoMS showed a similar correlation to treatment response, with higher numbers of presented neoantigens associated with clinical benefit (Fig.6B,E, Mann-Whitney test, p= 0.005 for the NSCLC cohort and p = 0.01 for the melanoma cohort). Interestingly, in the NSCLC cohort, response to treatment was not only associated with the number of mutations in the tumour, but also heavily positively correlated with the rate of presented neoantigens per mutation (Fig. 6C, Mann-Whitney test, p = 0.07): tumours for which ICI treatment led to durable clinical benefits had mutations that carried more presented neoantigens, It has to be noted that the latter was not due to the presence in ICI-responding tumours of a higher amount of frameshift or stop-loss mutations (hereafter named “high-yield mutations”) in ICI-responding tumours, inherently yielding more neoantigens and therefore likely to disproportionately affect the neoantigen presentation rate. Not only was the amount of presented neoantigens derived from high-yield mutations similar between responders and non-responders high-yield mutations actually generated more neoantigens in non-responders (Supp, Fig. 1). Furthermore, the rate of neoantigen presentation as predicted by neoMS allowed for further insights on treatment response prediction. Five of the NSCLC cohort members did not respond to ICI treatment despite a high TMB; as shown in Fig. 6C, the rate of neoantigen presentation of these five tumours actually fell well within the range of other non-responding tumours and was lower than both the responder median and the cohort median, potentially explaining the lack of success of ICI therapy in these cases. This observation was however specific to the cancer indication: there was no difference in neoantigen presentation rates between melanoma responders and non responders (Fig. 6F). Taken together, these results show that neoMS predictions of tumour-presented neoantigen landscapes correlate well with response to ICI treatment, and could be used in combination with other well-established metrics to refine predictions for response to ICI treatment in some indications.

**Fig. 6:**
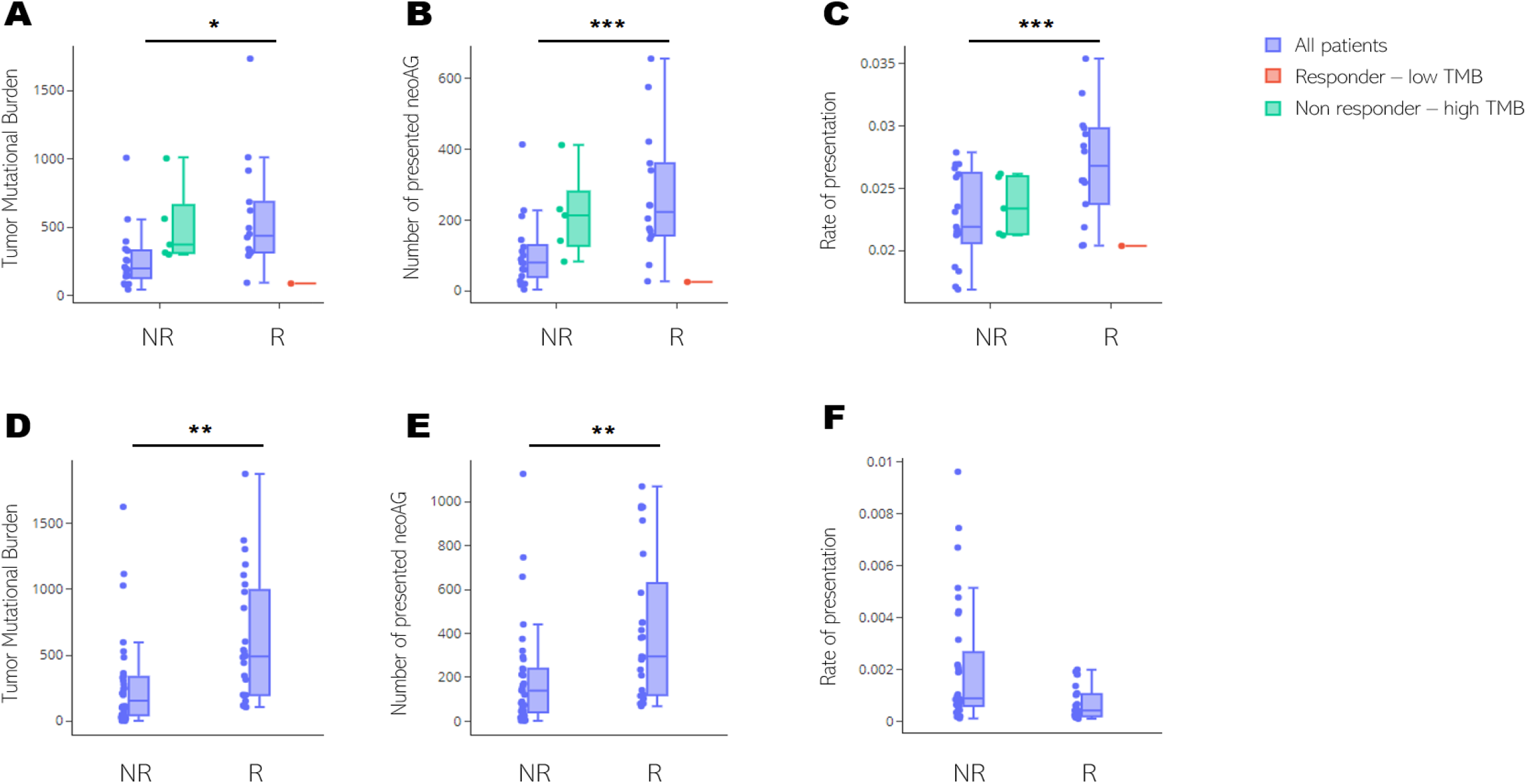
Mutational and neoantigen landscapes of tumors according to clinical benefit from CPI treatment in two cancer indications The tumor mutation burden (A,D), total number of presented neoantigens as predicted by neoMS (B,E) and rate of neoantigen presentation per mutation (C,F) are shown for a non-small cell lung cancer carcinoma (NSCLC) cohort (A-C) and a melanoma cohort (D-F). In the NSCLC cohort, patients with abnormal response relative to their TMB are also displayed with individual boxplots: non-responders with high TMB are shown in green and the responder with low TMB is shown in red. Asterisks mark significant differences between all responders and all non-responders according to Mann-Whitney tests: *: p<0.05, **: p = 0.01, ***: p<0.001

## Discussion

The neoMS algorithm shows unprecedented performance on MHC presentation prediction in clinically relevant samples. neoMS prediction scores also correlate well with peptide identification confidence levels in MS datasets, with high-confidence peptides more likely to be predicted as presented. Moreover, it has extrapolation capabilities, allowing good predictive powers even on HLA alleles not seen in training. These results make neoMS an attractive tool for any neoantigen discovery pipeline.

Through more accurate presentation predictions neoMS can improve neoantigen-driven therapy efficacy, due to the better target selection decreasing the amount of ineffective epitopes (*ie* false positives) used in therapy. Also, as not every tumour indication offers large enough biopsy amounts in clinical routine to perform MS-ligandome analysis on, the tool serves as a replacement of these extensive per-patient experiments. Furthermore, even in cases where MS ligandomes from tumors are obtained, an extra measure of confidence could be instated in the identification of these true positives, based on the a priori likelihood of presentation of the potential peptide on the cell surface estimated by a predictor such as neoMS. Overall, utilization of an accurate epitope presentation prediction tool holds the potential to greatly reduce therapy development costs and minimize therapeutic development timelines, by limiting the number of neoantigen validation experiments needed. Moreover, while additional training data would refine the neoMS model, its extrapolation capabilities allow neoMS to output reliable predictions with only minimal target supplementary data, Indeed, the ligandomic information of any HLA allele is able to inform prediction on all its close cognates, thus minimising the costly endeavor gathering additional MS-derived ligandome datasets.

Interestingly, neoMS predictions have been shown to correlate with response to ICI treatment in two cancer cohorts (NSCLC and melanoma) and can be used to refine predictions on the outcome of ICI treatment. Indeed, looking at the NSCLC cohort it was demonstrated that neoMS allows to further differentiate treatment response groups potentially improving treatment response predictions. In contrast, this could not be observed in the melanoma cohort, implying an indication-specific observation. We hypothesise this could likely be attributed to_differential response modalities to different types of immune checkpoint inhibitors (ipilimumab in the melanoma cohort and pembrolizumab in the NSCLC one) and/or intrinsic molecular differences between cancer types. Of note, more than 46% of the non-responders in the melanoma cohort (17/37) displayed a TMB higher than the best predictive cut point for response used in the NSCLC cohort (178 mutations), denoting strong inter-cohort differences (Rizvi et al., 2015; Snyder et al., 2014). Regardless of the cancer indication, however, the absolute number of presented neoantigens as predicted by neoMS is positively associated to clinical durable benefit with stronger statistical significance than TMB, supporting the biological relevance of the neoMS tool.

Some limitations however remain to the predictive power of the neoMS algorithm. As displayed by the model’s precision-recall curve and the sizable number of low-scoring peptides identified with high confidence in MS datasets, the number of false positives and negatives generated by the model remains non negligible. Further work will focus on improving the model performance, using several approaches. Custom amino-acid embedding using small value vectors derived from a principal component analysis of their main physico-chemical properties has shown to be effective in building high performance models and could improve prediction power by improving sequence representation. Previous work has also shown that the addition of peptide-extrinsic features (such as the flanking sequences present around the epitope in its protein of origin) can provide an increase in performance of up to 20%. Adding flanking sequences during the training of neoMS would allow the attention process to span them along with the epitope and HLA sequences, potentially yielding better predictions. In a third approach, a pretraining step using *masked language modeling* (*MLM*), where the model learns to fill in missing tokens in known input instances could improve sequence representation prior to presentation prediction training. In particular, MLM would allow us to leverage wide datasets of unlabeled data to learn representations that have been shown to be useful in later downstream tasks (Rives et al., 2000; Vig et al., 2020)

Finally, the neoMS architecture is theoretically not limited to MHC-I predictions for 9 to 11 mers. Provided that the reference sequences of the considered HLA alleles are known, and that some ligandomic data is available, the same framework could be used to infer MHC-II presentation likelihood. Although presentation mechanisms by HLA-II alleles are more complex, and diminished performance could therefore be expected, the neoMS algorithm can as such be extended into a MHC-II presentation predictor.

## Material and Methods

### Model structure

The neoMS algorithm follows a transformer, BERT-like architecture(Devlin et al., 2019), with 4 hidden encoder layers of dimensionality 72 and feed-forward layers of size 256. Each (self-)attention layers contains 6 attention heads, and the size of the vocabulary is set at 24 (special [CLS] token, separator [SEP] token, 20 amino-acid tokens and a unique “X” token for uncertain amino-acids). The output of the BERT part of neoMS is a tensor of the hidden states of the last layer for the first token of the sequence, further processed by a linear layer and a tanh activation function. On top of the BERT architecture is a 2-layer simple feed-forward network with input dimensionality 72, a hidden layer of size 256, and an output dimensionality of 1 with rectified linear activation function to process the results of the transformer network into a single probability.

### Positive instance selection

Positive instances were selected from previously published publicly available mass-spectrometry data(Abelin et al., 2017; Bulik-Sullivan et al., 2019; di Marco et al., 2017; Sarkizova et al., 2019; Trolle et al., 2016), that analysed the ligandome of both actual human tissue (with multiple HLAs) and cell lines expressing a single HLA allele. A peptide was considered a true positive if it was a 8-11mer detected by MS in a specific HLA allele (or HLA allele set) ligandome with a FDR of <0.02. The sequences of associated HLA alleles were obtained from the IPD-IMGT/HLA Database(Robinson et al., 2020). The reference sequences were processed such that for each HLA only the sequence in contact with the epitope is taken into account, in line with previous literature. Therefore, only the two alpha helices binding to the epitope were considered, i.e. residues 50-84 and 140-179, which will be subsequently called the “core sequence” in this text. A positive instance consisted of a tuple of the peptide sequence along with the core sequences of the entire set of HLA alleles (1 to 6) it was shown to be presented by, only considering alleles conclusively identified in the study of origin. Duplicates were dropped from the positive dataset: if a peptide was detected several times by the same set of alleles, it only accounted for a single positive instance. Detected peptides exhibiting non canonical amino acids were likewise discarded. Positive data after such filtering steps comprised 386647 instances, of which 173361 were derived from multiallelic HLA sets and 213286 from monoallelic HLAs. The monoallelic training dataset included ligandomic information from 92 different HLA alleles, while 76 different HLA alleles were present within the multiallelic dataset across 66 unique combinations; Table 1 summarises the characteristics of the entire positive dataset.

**Table 1:** Characteristics of the positive dataset

|  | Number of ligandomes | Number of individual HLA alleles | Number of instances |
| --- | --- | --- | --- |
| Multiallelic data | 66 | 76 | 173,361 |
| Monoallelic data | 92 | 92 | 213,286 |
| **TOTAL** | **158** |  | **386,647** |

### Train, validation and test datasets construction

Datasets were constructed as described previously(Bulik-Sullivan et al., 2019), and are summarised in Table 2. Briefly, two test datasets were constructed and held out from training and validation data: (i) a multiallelic sample test set consisting of 729 presented peptides from five tumour samples, and (ii) a monoallelic sample test consisting of 2039 presented peptides from a cell line expressing the A*74:01 allele. The first dataset aims at evaluating the performance of the model on actual novel patient samples; it is also identical to the test dataset used by Bulik-Sullivan et al. to allow for comparison between the two algorithms (Bulik-Sullivan et al., 2019). The second dataset aims at testing the extrapolation capabilities of the model on previously unseen HLA alleles: the A*74:01 allele is not present in any of the other training instances, and its core sequence is different from any other core sequence present in the train dataset. For these test datasets, negative instances were synthesised at a ratio of 1 positive instance for 2,500 negative instances (1:2500).

**Table 2:**
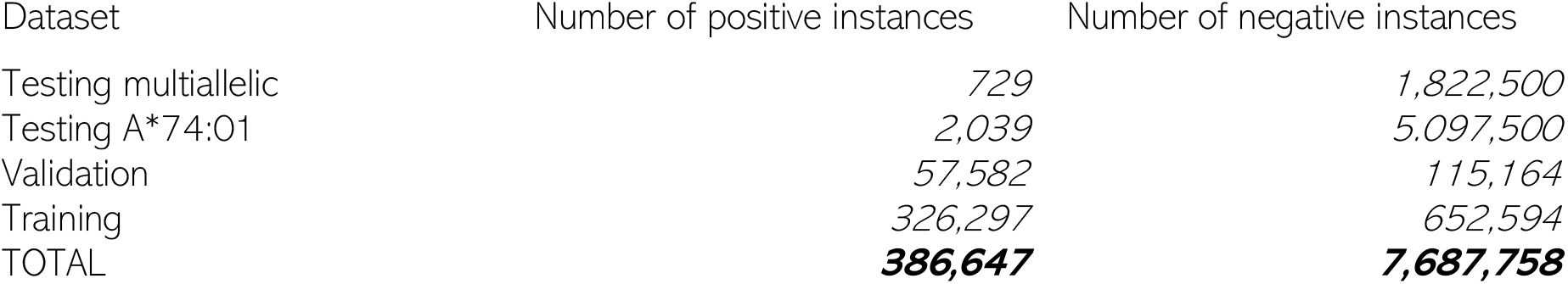
Characteristics of the training, testing and validation datasets

| Dataset | Number of positive instances | Number of negative instances |
| --- | --- | --- |
| Testing multiallelic | 729 | 1,822,500 |
| Testing A*74:01 | 2,039 | 5,097,500 |
| Validation | 57,582 | 115,164 |
| Training | 326,297 | 652,594 |
| TOTAL | **386,647** | **7,687,758** |

The remaining 383879 positive instances were randomly split into a training set (85% of the data) and a validation set (15%). For these datasets, negative instances were synthesised at a ratio of 1 positive instance for 2 negative instances in order to preserve class balance(1:2). The validation dataset was only used for evaluation of model performance during training and early stopping.

Since positive data was deduplicated, no positive instances could be present in several datasets. In addition, negative instances were screened so that no negative instance appears in more than one dataset; when a negative instance was detected in several datasets, all its occurrences but one were discarded and new negative instances were synthesised in their place.

### Negative instance synthesis

Negative peptide instances were created *in silico* using the following procedure: first, a single or set of HLA was selected at random from the pool of positive instances from the considered dataset. Since this choice was weighted by the frequency of occurrence of a given HLA set in the positive instances, the distribution of HLAs of the negative set matched the distribution of the positive set. Then, a 8-11mer was selected at random from the reference human proteome (EMBL-EBI, v.96); it was ensured that the negative peptide was not present in association with the selected HLA set in any of the datasets. It should be noted it is possible that some of the peptides labeled as negative using this strategy could actually be presented in certain conditions or indications; however, the low rate of peptide presentation means that such false negatives would be rare in a randomly sampled dataset. The negative instance consisted of a tuple of the sequence of the randomly sampled 8-11mer along with the core sequences of the selected HLA(s). To further ensure separation between negative instances of the train/validation datasets on the one hand and the test dataset on the other hand, the human reference was randomly split into two sets of proteins. Negative instances from the test datasets were sampled from one of the sets whereas negative instances from the train and validation datasets were sampled from the other.

### Peptide and HLA encoding for machine learning

For each instance, peptide and HLA sequences were encoded as tokens, wherein each amino-acid from each sequence was encoded with its specific token. All peptides were padded to length 11 using pad tokens. To allow the model to focus on relevant amino-acids, all HLA core sequences were aligned to the core sequence of A*02:52; subsequently, only amino acids differing from this reference core sequence were encoded, the rest was replaced by pad tokens. HLA core sequences of the instance were delimited with specific tokens at the start of the sequence of each helix, after which they were concatenated into a single vector. Finally, a 87- to 379-long instance tensor was constructed initiated by a CLS token, and comprised of the peptide and HLA(s) encodings separated by SEP tokens. The two parts of the core sequences of each HLA allele in the instance (*ie* the two alpha helices) were prefaced by specific tokens, denoted [HLA-1] and [HLA-2]. Correct attribution of tokens to peptide or HLA was provided to the model using a typing mask.

#### Training

Instances were considered as independent, and the per-peptide loss was defined as the binary cross-entropy loss function. Formally, the contribution of instance *i* to the overall loss is

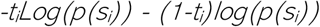

Where *t_i_* is the ground truth label of instance *i* (1 for a peptide presented by the HLA set and 0 for a peptide not presented) and *p*(*t_i_*) is the predicted probability for the instance to belong to its ground truth class. Model weights were initialised following a truncated normal distribution of standard deviation 0.02 and trained using the ADAM optimizer (Kingma & Ba, 2015)with default parameters on an Amazon p2 instance using Nvidia K80 GPUs. The validation set was used for monitoring and early stopping of the training procedure. Instances were shuffled and processed using a batch size of 96. The learning rate was increased every 100 batches of the first 2 epochs using a gradual warmup scheduler, after which it was decreased by half every other epoch (https://github.com/ildoonet/pytorch-gradual-warmup-lr).

### Performance metrics and precision recall curves

Precision and recall were chosen as performance metrics. With true positives, false positives and false negatives identified respectively as TP, FP and FN, precision (also known as Positive Predictive Value or PPV) is defined as:

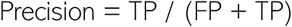

And recall as:

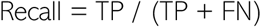

Epitope presentation is a problem with highly imbalanced classes, with most possible epitopes not being presented at the surface of the cell. Moreover, positive events are highly desirable, as potentially actionable epitopes solely fall within that category. As such, precision and recall are a good assessment of an epitope presentation prediction model, reflecting both its ability to capture as much of the available true positive pool as possible and its ability to minimise false positives.

Precision and recall for neoMS were calculated by confronting the results of the model on the test data to the ground truth using the Python *scikit-learn* library. For the EDGE model, precision and recall values were obtained for the MS-only model by averaging the values obtained for each tumour sample contained in the test dataset, as displayed in the original paper.

### MHCflurry

Binding affinities of peptides in the test set were predicted using *MHCflurry* 1.6.1 (O’Donnell et al., 2018). *MHCflurry* is an open-source MHC-I binding affinity predictor with performance comparable or superior to the *netMHC* family of models, and therefore representative of the best affinity predictors (Zhao & Sher, 2018). To combine binding affinity predictions for a single peptide across a set of HLAs (ie an actual test set instance), the minimal affinity prediction was chosen for each peptide. Precision and recall were computed using this approach on the test dataset.

### Motif logos

Motif logos were generated using the weblogolib Python API(Schneider & Stephens, 1990). Model predictions were calculated for 2,000,000 random peptides for each peptide length considered. The top 1% predictions were used to generate the logo.

### Attention

Attention processes in the neoMS model were generated using *bertviz head view* (https://github.com/jessevig/bertviz).

### Ligandome analysis

MS data obtained from HLA-I immunoprecipitates was obtained from Loffler et al, (Löffler et al., 2019), converted to mgf format using MSConvert(Chambers et al., 2012) and peptide identification was performed using Comet (Eng et al., 2013) and MS-GF+ (Kim & Pevzner, 2014) provided by SearchGUI 3.3.21 (Vaudel et al., 2011) at a PSM FDR threshold of 1%. Oxidation of methionine and N-terminal acetylation were specified as variable modifications. The maximum allowed precursor m/z tolerance was set to 10 ppm, and the fragment m/z tolerance to 0.5 Da. The PSMs were further analysed for peptide and protein matches using PeptideShaker 1.16.45, giving a per peptide confidence score (Vaudel et al., 2015). neoMS predictions were then computed for every 8-11 mer identified using this procedure.

### Cancer cohort analyses

Mutation lists, HLA types and response to treatment information were obtained from publicly available cohorts(Rizvi et al., 2015; Snyder et al., 2014). The exhaustive set of mutation-derived 8-11mer neoantigens was identified for each sample using a modified version of MuPeXI (Bjerregaard et al., 2017), and used as input for neoMS

