## Supplemental Figure 1 for "neoMS: Attention-based Prediction of MHC-I Epitope Presentation"

**Supplementary figures**

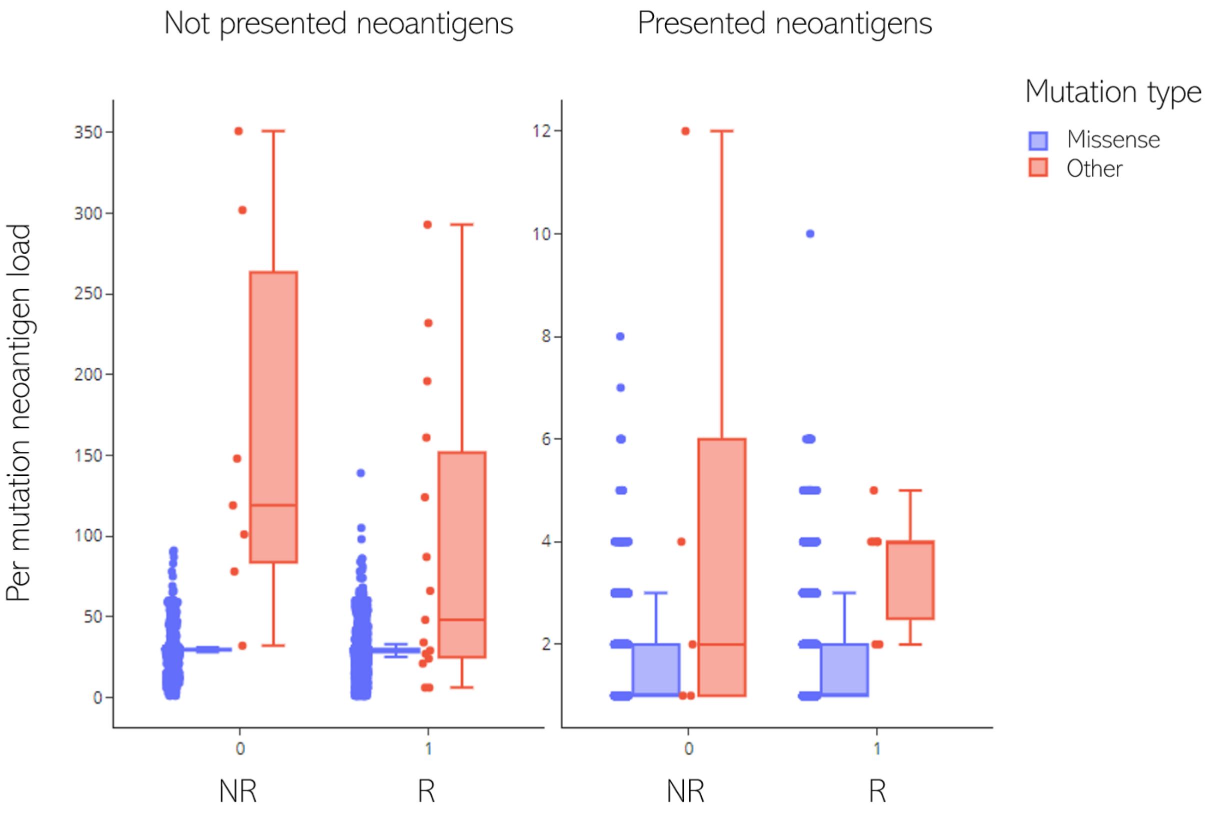


*Supp. Fig. 1: Neoantigen load per mutation type in the lung cancer patient cohort*

*Each data point represents the amount of non-presented (left) and presented (right) neoantigens derived from a given mutation. "Other" mutations can include in-frame insertions, deletions, frameshifts and stop-loss. In the cohort considered, only stop-loss mutations were detected. R: Non Responders, R: Responders*
